# Putrescine-responsive post-transcriptional feedback regulation of Arabidopsis *ADC2* mediated by 5’-UTR elements

**DOI:** 10.64898/2026.10.31.763274

**Authors:** Yuta Hiragori, Miharu Yasumuro, Taihei Karino, Atsushi Kaido, Yasuko Sakihama, Yuya Goto, Yui Yamashita, Satoshi Naito, Hitoshi Onouchi

**Affiliations:** Graduate School of Agriculture, Hokkaido University, Sapporo 060-8589, Japan; Research Faculty of Agriculture, Hokkaido University, Sapporo 060-8589, Japan; School of Agriculture, Hokkaido University, Sapporo 060-8589, Japan

## Abstract

Putrescine (Put) is a bioactive polyamine that contributes to plant growth, development, and stress responses, whereas excessive Put accumulation impairs growth. However, how plants maintain Put levels within an appropriate range remains poorly understood. In *Arabidopsis thaliana*, expression of the arginine decarboxylase 2 (*ADC2*) gene is a major determinant of Put abundance. Here, we show that the 5’ untranslated region (5’-UTR) of *ADC2* mediates Put-responsive feedback regulation and contributes to Put homeostasis in Arabidopsis. In transient expression assays using an emerald luciferase (Eluc) gene placed downstream of the *ADC2* 5’-UTR (*ADC2* 5’-UTR:*Eluc*), Eluc activity was reduced by exogenously applied Put but increased when endogenous Put biosynthesis was impaired by *adc2* mutations or treatment with an arginine decarboxylase inhibitor. In transgenic plants carrying *ADC2* 5’-UTR:*Eluc*, Put treatment modestly decreased *Eluc* mRNA abundance but substantially reduced Eluc activity and translation efficiency, suggesting that this Put-responsive repression occurs mainly at the translational level. Among the *cis*-regulatory elements in the *ADC2* 5’-UTR, mutations affecting either the conserved AUU/CUG-initiated upstream ORF or the predicted RNA secondary structure in the ADC-box, a conserved GC-rich motif, abolished Put-responsive repression, indicating that these elements are required for this regulation. Two genome-edited lines carrying deletions spanning these elements exhibited delayed germination and impaired growth, and the more severely affected line showed increased Put accumulation. Together, these findings reveal a feedback mechanism that regulates *ADC2* expression through its 5’-UTR and support a role for this mechanism in Put homeostasis in Arabidopsis.

## INTRODUCTION

Polyamines are essential and ubiquitous molecules in living cells. Owing to their cationic nature, polyamines interact with DNA, RNA, ATP, and other negatively charged cellular molecules, thereby influencing multiple stages of gene expression, including translation (Kusano et al. 2008; Igarashi and Kashiwagi 2010). The major polyamines in many species are the diamine putrescine (Put), the triamine spermidine (Spd), and the tetraamine spermine (Spm) (Fig. 1). Plants also produce thermospermine (Tspm), a structural isomer of Spm (Blázquez 2024). At appropriate levels, polyamines are required for normal plant growth and development, and contribute to tolerance to biotic and abiotic stresses (Chen et al. 2019; Blázquez 2024). By contrast, excessive accumulation of polyamines has adverse effects. For example, transgenic tobacco and Arabidopsis lines overexpressing Put biosynthetic genes exhibit Put overaccumulation and severe growth defects, such as leaf chlorosis and necrosis in tobacco and dwarfism and late flowering in Arabidopsis (Masgrau et al. 1997; Alcázar et al. 2005). The dual nature of Put suggests the presence of mechanisms that tightly maintain its homeostasis, yet these mechanisms remain poorly understood in plants.

**Figure 1.**
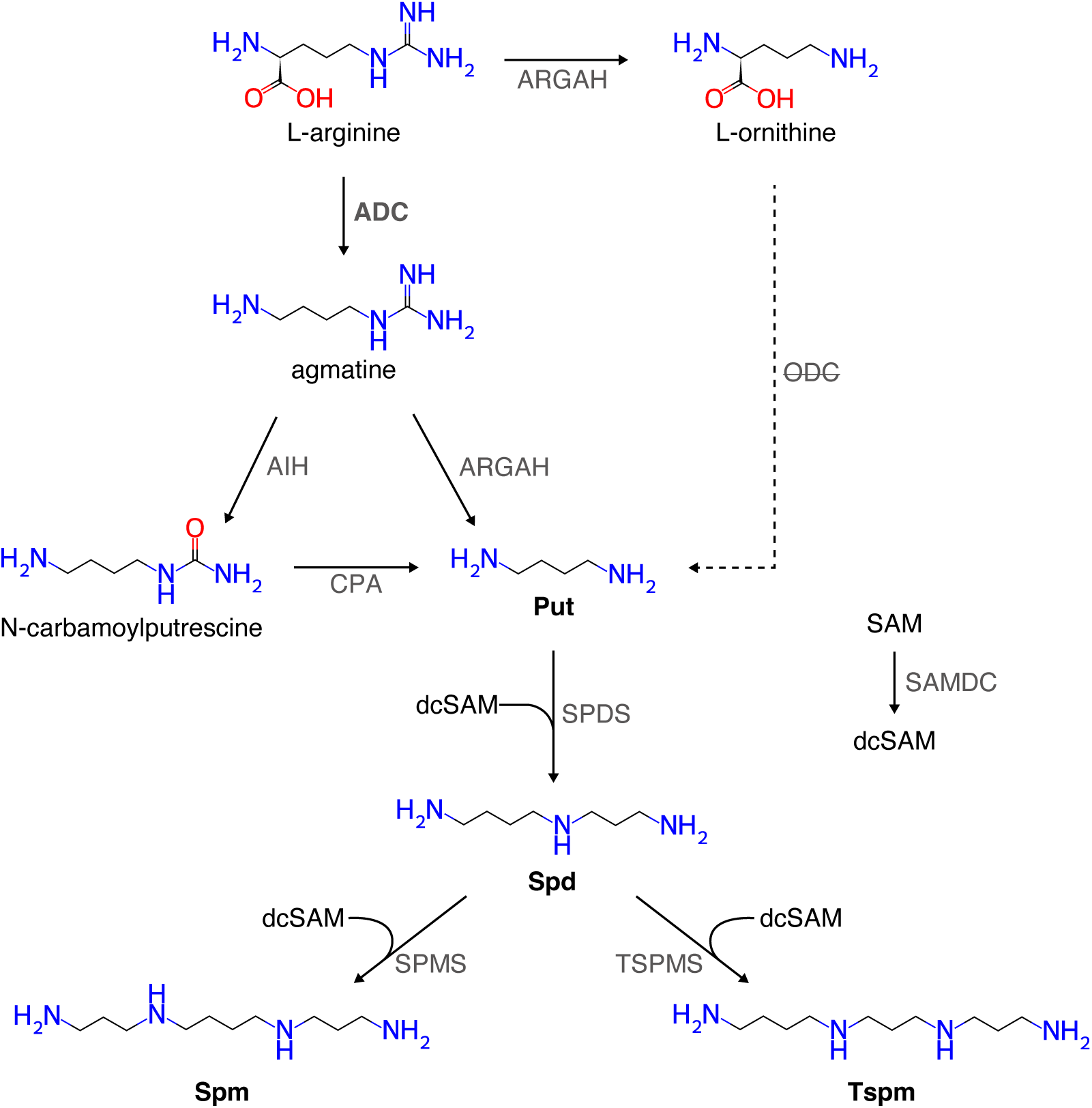
Polyamine biosynthetic pathways relevant to Put homeostasis in Arabidopsis. Put is produced from Arg through arginine decarboxylase (ADC), followed by agmatine iminohydrolase (AIH) and *N*-carbamoylputrescine amidohydrolase (CPA), or by the agmatinase activity of arginase (ARGAH). ARGAH also converts Arg to Orn. The dashed ornithine decarboxylase (ODC) route is shown for comparison; Arabidopsis lacks a functional ODC pathway. Put is converted to Spd, which is used to synthesize Spm or Tspm. SPDS, spermidine synthase; SPMS, spermine synthase; TSPMS, thermospermine synthase. *S*-adenosylmethionine decarboxylase (SAMDC) generates decarboxylated *S*-adenosylmethionine (dcSAM), the aminopropyl donor, from *S*-adenosylmethionine (SAM).

Put occupies a central position in plant polyamine biosynthesis as the precursor to other polyamines, Spd, Spm, and Tspm (Fig. 1). In many plant species, Put is synthesized either from L-arginine (Arg) through a pathway initiated by arginine decarboxylase (ADC) or directly from L-ornithine (Orn) by ornithine decarboxylase (ODC). In the ADC pathway, ADC first converts Arg to agmatine. Agmatine is then converted to Put either via *N*-carbamoylputrescine (Janowitz et al. 2003; Piotrowski et al. 2003) or directly by arginases with agmatinase activity as reported in Arabidopsis and soybean (Patel et al. 2017). In the ODC pathway, ODC directly converts Orn to Put (Michael et al. 1996). In *Arabidopsis thaliana*, however, the genome lacks a functional *ODC* gene, and Put biosynthesis therefore depends solely on the ADC pathway (Hanfrey et al. 2001; Fig. 1). The Arabidopsis genome contains two genes encoding ADC, *ADC1* (At2g16500, also called *SPE1*) and *ADC2* (At4g34710, also called *SPE2*). Histochemical analyses using β-glucuronidase staining showed that *ADC1* promoter activity was generally low during vegetative development, whereas the *ADC2* promoter was active during seed germination and root and leaf development (Hummel et al. 2004). Moreover, loss of *ADC2* significantly reduces Put levels, whereas loss of *ADC1* does not (Urano et al. 2004; Rossi et al. 2015), supporting a predominant role for *ADC2* in determining Put levels in Arabidopsis.

Feedback regulation of polyamine biosynthetic genes contributes to polyamine homeostasis in eukaryotes. A well-characterized example is *S*-adenosylmethionine decarboxylase (SAMDC), which produces decarboxylated *S*-adenosylmethionine, an aminopropyl donor required for Spd, Spm, and Tspm synthesis (Fig. 1). *SAMDC* mRNAs contain an upstream open reading frame (uORF) with an evolutionarily conserved amino acid sequence (Franceschetti et al. 2001). uORFs can repress translation of the downstream main ORF by causing ribosome dissociation after uORF translation or by inducing ribosome stalling during uORF translation (von Arnim et al. 2014; Hinnebusch et al. 2016; Dever et al. 2020). In some cases, they can also trigger mRNA decay through quality-control pathways such as nonsense-mediated decay (NMD) (Gaba et al. 2005; Nyikó et al. 2009; Uchiyama-Kadokura et al. 2014). In mammals and plants, elevated Spd/Spm levels induce ribosome stalling during translation of the *SAMDC* uORFs, thereby repressing *SAMDC* translation. This ribosome stalling depends on the uORF-encoded peptide, although the conserved peptide sequences differ between mammals and plants (Ruan et al. 1996; Law et al. 2001; Raney et al. 2002; Hanfrey et al. 2002; Hanfrey et al. 2005; Uchiyama-Kadokura et al. 2014). In addition to this translational feedback regulation, *ACAULIS5* (*ACL5*) mRNA, which encodes Tspm synthase, is subject to transcriptional feedback regulation by Tspm (Kakehi et al. 2008; Yariuchi et al. 2021). However, feedback regulation of genes encoding Put biosynthetic enzymes has not been established in plants.

Previous studies have identified several *cis*-regulatory elements in the 5’ untranslated regions (5’-UTRs) of plant *ADC* genes that repress expression of the downstream main ORF. Earlier work showed that a short uORF in the carnation *ADC* gene represses translation of the main ORF (Chang et al. 2000). More recently, a conserved GC-rich motif termed the “ADC-box” was identified in the 5’-UTRs of plant *ADC* genes and shown to repress translation of the main ORF in tomato. The ADC-box was predicted to form a stable RNA secondary structure that may contribute to repression by impeding ribosome scanning (Wu et al. 2019). Additionally, we recently identified a conserved non-AUG-initiated uORF in plant *ADC* genes and demonstrated its repressive effect in Arabidopsis *ADC2*. In the *ADC2* uORF, two noncanonical start codons, AUU and CUG, contribute to repression of the main ORF (Hiragori et al. 2023). Notably, the 5’-UTR of Arabidopsis *ADC2* contains other putative *cis*-regulatory elements, including a short AUG-initiated uORF and the ADC-box. However, the physiological significance of the cis-regulatory elements in the *ADC2* 5’-UTR remains unclear.

Here, we investigated whether the Arabidopsis *ADC2* 5’-UTR mediates Put-responsive regulation and contributes to Put homeostasis. Reporter assays showed that the *ADC2* 5’-UTR represses main ORF expression in response to exogenous Put, whereas inhibition of endogenous Put biosynthesis has the opposite effect. Analyses of transgenic reporter lines further indicated that translational repression makes a major contribution to this response. Mutational analyses revealed that Put-responsive repression requires two elements within the *ADC2* 5’-UTR: the AUU/CUG-initiated uORF and the predicted RNA secondary structure of the ADC-box. Genome-edited *ADC2* 5’-UTR mutants exhibited delayed germination and impaired growth, and the more severely affected mutant overaccumulated Put. Together, these findings reveal a Put-responsive feedback mechanism mediated by the *ADC2* 5’-UTR and support a role for the regulatory functions of this region in maintaining Put homeostasis and normal growth in Arabidopsis.

## RESULTS

### The Arabidopsis *ADC2* 5’-UTR regulates downstream main ORF expression in response to intracellular Put levels

To examine whether the Arabidopsis *ADC2* 5’-UTR mediates repression of downstream main ORF expression in response to Put, we performed transient expression assays in Arabidopsis mesophyll protoplasts. The *ADC2* 5’-UTR and the first 5 nt of its main ORF were fused to the coding sequence of emerald luciferase (Eluc) carrying a PEST degradation sequence (*Eluc*-PEST), under the control of the cauliflower mosaic virus 35S RNA promoter (35Spro). This construct was designated as *ADC2* 5’-UTR:*Eluc*. A control reporter carried only a short unrelated sequence in place of the *ADC2* 5’-UTR (Fig. 2A). A plasmid expressing *Renilla* luciferase (Rluc) from the same promoter was co-introduced, and Eluc activity was normalized to Rluc activity. After 24 h of incubation with 0–4 mM Put, normalized Eluc activity from *ADC2* 5’-UTR:*Eluc* showed a strong negative correlation with exogenous Put concentration (Fig. 2B). The control construct produced substantially higher Eluc activity than *ADC2* 5’-UTR:*Eluc* and showed no significant response to 2 mM Put, whereas activity from *ADC2* 5’-UTR:*Eluc* was reduced by this treatment (Fig. 2C). The absence of a significant response in the control construct indicates that the observed Put-responsive repression was not a general effect of Put on the 35Spro or the *Eluc*-PEST reporter system. Thus, the *ADC2* 5’-UTR conferred both strong basal repression and repression in response to exogenous Put.

**Figure 2.**
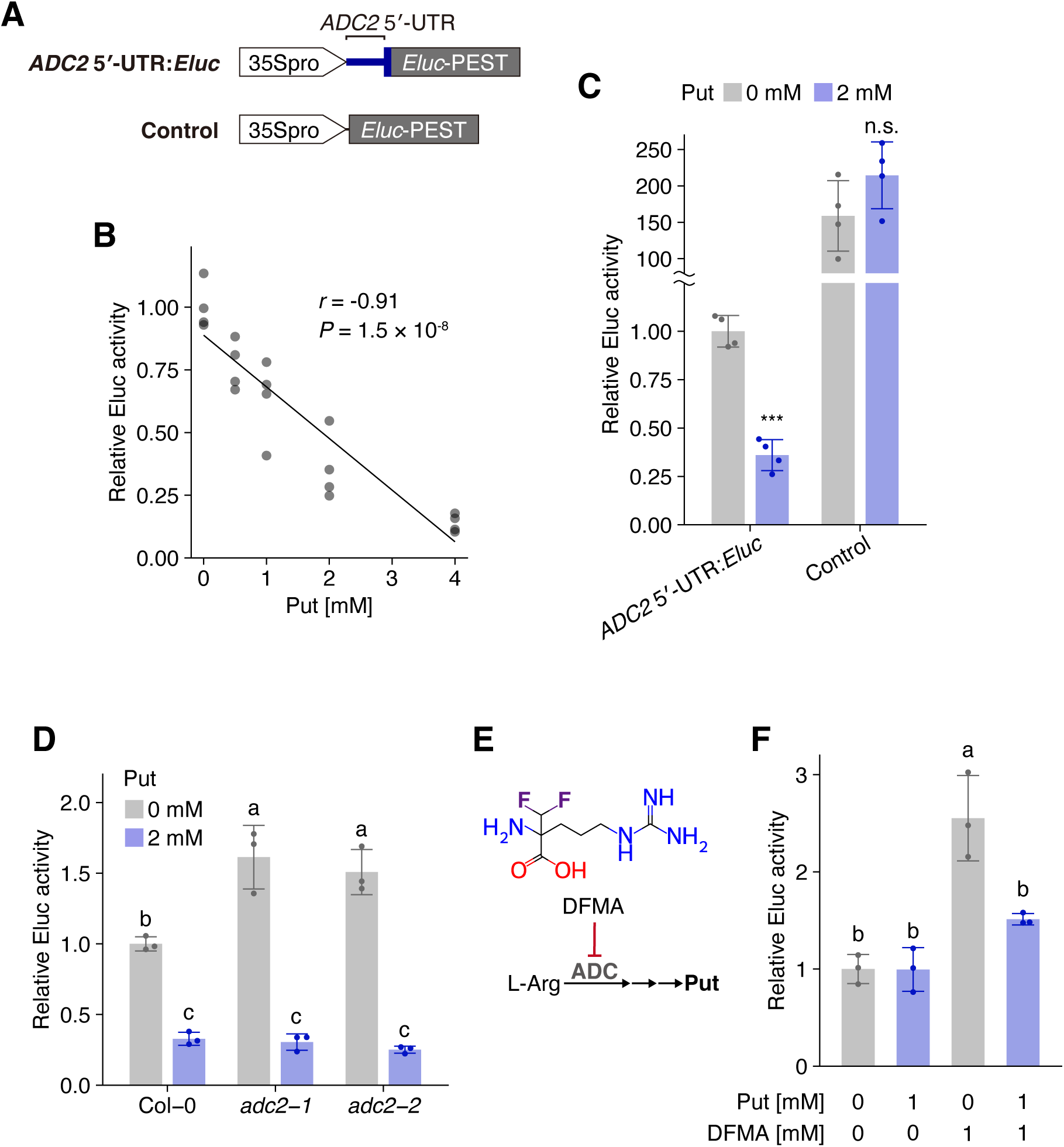
The *ADC2* 5’-UTR mediates reporter responses to Put supply and inhibition of Put biosynthesis. (A) Schematic representations of reporter constructs. *ADC2* 5’-UTR:*Eluc* (upper) contains the *ADC2* 5’-UTR (thick blue line) and the first 5 nt of its main ORF (blue box) followed by *Eluc*-PEST (gray box). The control construct (lower) contains a short unrelated sequence in place of the *ADC2*-derived sequence. Both reporters are driven by 35Spro (white arrows). (B) Relative Eluc activity in mesophyll protoplasts carrying the *ADC2* 5’-UTR:*Eluc* plasmid after 24 h of treatment with 0–4 mM Put (n = 4 per concentration). The line was fitted by least-squares regression; Pearson’s correlation coefficient (*r*) and *P* value are *r* = −O.91 and *P* = 1.5 × 1Ō8, respectively. (C) Relative Eluc activity from the *ADC2* 5’-UTR:*Eluc* and control plasmids in mesophyll protoplasts treated with 0 or 2 mM Put (n = 4 per group). Note the discontinuous y-axis. (D) Relative activity of the *ADC2* 5’-UTR reporter in mesophyll protoplasts from Col-0, *adc2-1*, and *adc2-2*, with 0 or 2 mM Put (n = 3 per group). (E) Chemical structure of DFMA and schematic inhibition of ADC-dependent Put biosynthesis. (F) Relative activity of the *ADC2* 5’-UTR reporter in MM2d suspension cell protoplasts treated with 0 or 1 mM Put and 0 or 1 mM DFMA (n = 3 per group). In all transient assays, protoplasts were incubated at 22°C for 24 h in the dark. Eluc activity was normalized to Rluc activity from the reference plasmid and scaled to the mean of untreated *ADC2* reporter control in B and C, untreated Col-0 in D, or the no-addition control in F, within each experiment. Bars show means ± SD. In C, data were compared between the 0 and 2 mM Put treatments within each construct using one-sided, unpaired Student’s *t* tests assuming equal variances, with Holm correction for the two comparisons (***, adjusted *P* < 0.001; n.s., not significant, adjusted *P* ≥ O.O5). In D and F, different letters indicate significant differences among groups within each panel (two-way ANOVA followed by Tukey’s test, *P* < 0.05).

We next tested whether perturbation of endogenous Put biosynthesis affected Eluc activity from the *ADC2* 5’-UTR reporter. We first used protoplasts prepared from two *adc2* mutants carrying T-DNA insertions in *ADC2*, *adc2-1* (SALK_073977) and *adc2-2* (WiscDsLoxHs065_09H; Supplemental Fig. S1A). The *adc2-1* mutant has been reported to reduce endogenous *ADC2* expression and Put levels (Kim et al. 2013). Reverse transcription quantitative PCR (RT-qPCR) analysis showed that *ADC2* transcript abundance was markedly reduced in both mutants compared with Columbia-0 (Col-0) wild-type (WT), with only one of three biological replicates of *adc2-1* and none of those of *adc2-2* yielding quantifiable values (Supplemental Fig. S1B), suggesting that endogenous Put levels may also be reduced in *adc2-2*. In protoplasts prepared from both mutants, relative Eluc activity from *ADC2* 5’-UTR:*Eluc* was higher than that in Col-0 in the absence of exogenous Put. After treatment with 2 mM Put, relative Eluc activity was reduced in all genotypes, and the differences observed in untreated conditions were no longer detected (Fig. 2D).

We further examined the effect of inhibiting endogenous Put biosynthesis in protoplasts prepared from Arabidopsis MM2d suspension cells (Menges and Murray 2002). Protoplasts transfected with *ADC2* 5’-UTR:*Eluc* were incubated with or without 1 mM a-difluoromethylarginine (DFMA), an ADC inhibitor reported to reduce endogenous Put levels (Birecka et al. 1985; Fallon and Phillips 1988; Fig. 2E). Unlike in mesophyll protoplasts, 1 mM Put alone did not significantly reduce relative Eluc activity in MM2d protoplasts (Fig. 2, B and F). Because increases in Put levels have been reported during callus induction and cellular reprogramming (Desai and Mehta 1985; Iwase et al. 2018), this difference may reflect differences in endogenous Put levels associated with the distinct differentiation states of the source cells. DFMA alone significantly increased Eluc activity, whereas addition of 1 mM Put in the presence of DFMA significantly reduced it (Fig. 2F). These results support the interpretation that the DFMA-induced increase in Eluc activity resulted from a decrease in endogenous Put levels. Together, the responses to exogenous Put, *adc2* mutations, and ADC inhibition demonstrate feedback regulation mediated by the *ADC2* 5’-UTR in response to intracellular Put levels.

### Put-responsive repression mediated by the *ADC2* 5’-UTR involves regulation at both the mRNA and translational levels

We next examined the regulatory level at which the *ADC2* 5’-UTR mediates Put-responsive repression. Changes in luciferase activity alone cannot distinguish altered translation from altered reporter mRNA abundance. We therefore generated Arabidopsis transgenic lines carrying the *ADC2* 5’-UTR:*Eluc* construct and conducted reporter assays together with RT-qPCR. Two independent homozygous *ADC2* 5’-UTR:*Eluc* lines were analyzed. As a negative control, we used a homozygous line carrying the 5’-UTR of *MASP1* (At4g03260) in the same reporter vector (Fig. 3A). Ten-day-old seedlings were incubated with or without 2 mM Put for 24 h, after which Eluc activity and *Eluc* mRNA abundance were measured. Put treatment significantly reduced Eluc activity in both independent *ADC2* 5’-UTR:*Eluc* lines to approximately 40% of their respective untreated levels but had no significant effect on Eluc activity in the *MASP1* control line (Fig. 3B).

**Figure 3.**
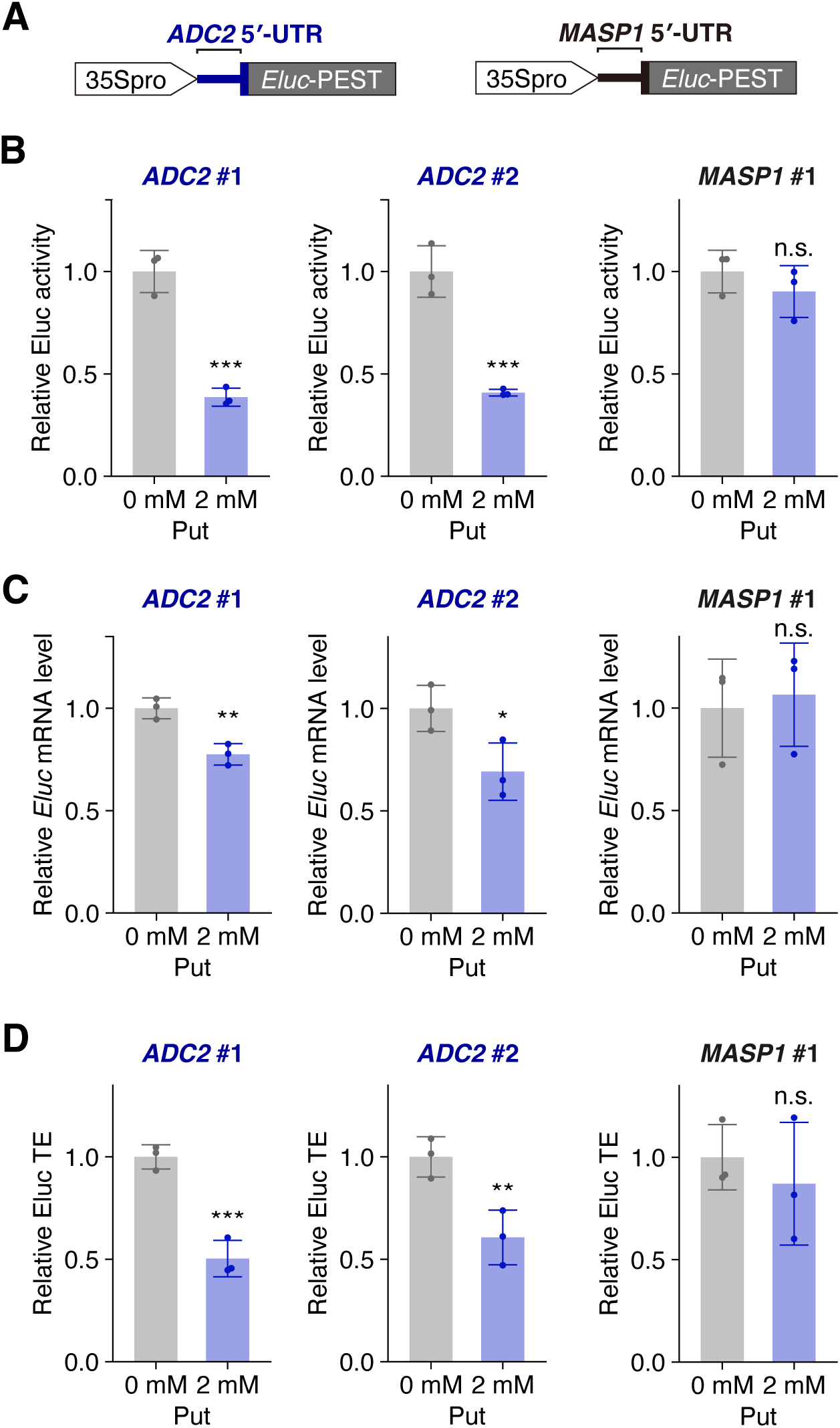
Put reduces reporter activity, mRNA abundance, and TE in stable *ADC2* 5’-UTR reporter lines. (A) Schematic representations of the reporter constructs used for transgenic reporter assays. These constructs contain the 5’-UTR and the first 5 nt of the main ORF from either *ADC2* (blue thick line and box) or *MASP1* (black thick line and box) upstream of *Eluc*-PEST (gray box), driven by the 35Spro (white arrow). (B) Relative Eluc activity in two independent *ADC2* reporter lines (*ADC2* #1 and *ADC2* #2) and the *MASP1* control line (*MASP1* #1). Ten-day-old seedlings were transferred to liquid half-strength MS medium and treated with 0 or 2 mM Put for 24 h. Eluc activity was normalized to total protein. (C) Relative *Eluc* mRNA abundance, normalized to *EF1a* mRNA in the same cDNA sample. (D) Relative Eluc TE, calculated for each biological replicate as relative Eluc activity divided by relative *Eluc* mRNA abundance measured in the same sample. Values are expressed relative to the untreated condition for each line. Bars show means ± SD of three biological replicates, each comprising 15 seedlings pooled from one flask. Treated and untreated samples were compared within each line using one-sided, unpaired Student’s *t* tests assuming equal variances. *, *P* < 0.05; **, *P* < 0.01; ***, *P* < 0.001; n.s., *P* ≥ 0.05.

Put treatment also caused a modest but statistically significant decrease in *Eluc* mRNA abundance in both *ADC2* 5’-UTR:*Eluc* lines, to approximately 70% to 80% of their respective untreated levels, whereas no significant change was detected in the *MASP1* control line (Fig. 3C).

We calculated relative Eluc translational efficiency (TE) as the ratio of relative Eluc activity to relative *Eluc* mRNA abundance. Put treatment significantly reduced relative Eluc TE in both *ADC2* 5’-UTR:*Eluc* lines to approximately 50% to 60% of their respective untreated levels, whereas no significant change was detected in the *MASP1* control line (Fig. 3D). The reduction in relative Eluc TE (approximately 40%–50%) was greater than the reduction in reporter mRNA abundance (approximately 20%–30%). These results support translational repression as a major component of the Put-responsive regulation mediated by the *ADC2* 5’-UTR, with a smaller contribution from reduced reporter mRNA abundance.

### The non-AUG-initiated uORF is required for Put-responsive regulation mediated by the *ADC2* 5’-UTR

We next determined which *cis*-elements within the *ADC2* 5’-UTR are required for Put-responsive regulation. A short AUG-initiated uORF and a conserved non-AUG-initiated uORF have been identified in plant *ADC* 5’-UTRs (Chang et al. 2000; Hiragori et al. 2023), and both uORFs are present in the Arabidopsis *ADC2* 5’-UTR (Fig. 4A; Supplemental Fig. S2). The conserved ADC-box motif is also present in the *ADC2* 5’-UTR and overlaps with the AUU/CUG-initiated uORF (Fig. 4A; Supplemental Fig. S2).

**Figure 4.**
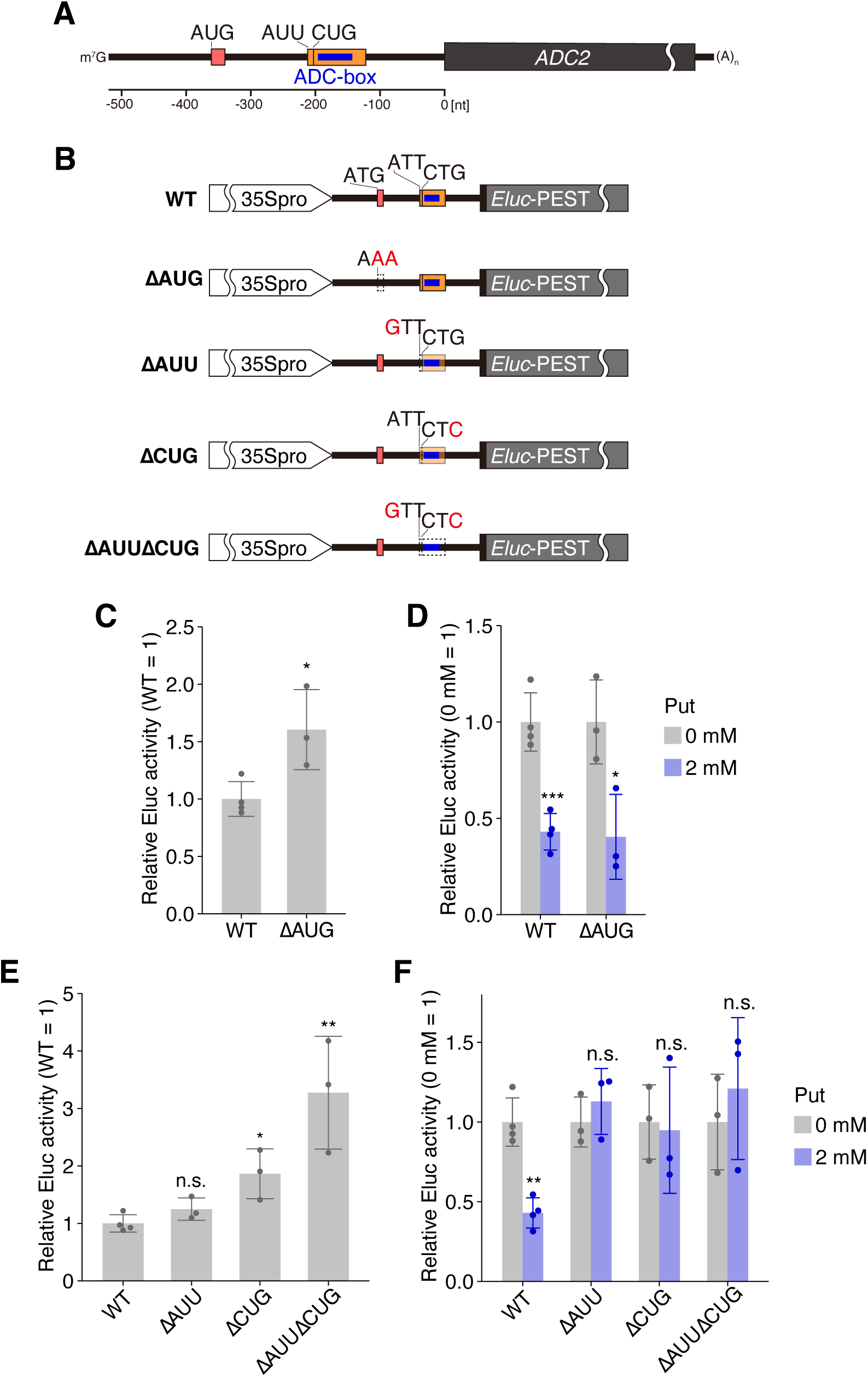
Mutational analysis of the AUG- and non-AUG-initiated uORFs in the *ADC2* 5’-UTR. (A) Organization of the *ADC2* 5’-UTR. The short AUG-initiated uORF is shown in red, the AUU/CUG-initiated uORF in orange, and the ADC-box in blue. Positions are shown relative to the main ORF start. mၭG, 5’ cap; (A)n, poly(A) tail. (B) WT, ΔAUG, ΔAUU, ΔCUG, and ΔAUUΔCUG reporter constructs. The DNA start codon ATG was changed to AAA in ΔAUG; ATT was changed to GTT and CTG to CTC in the non-AUG mutants, individually or together. Red letters indicate substitutions. Schematics use DNA notation, whereas construct names refer to RNA codons. (C, E) Relative Eluc activity in untreated mesophyll protoplasts, with WT set to 1, for ΔAUG (C) and the non-AUG mutants (E). (D, F) Relative Eluc activity after treatment with 0 or 2 mM Put, with the untreated value set to 1 for each construct. Assays were incubated at 22°C for 24 h in the dark. Eluc activity was normalized to Rluc activity from the reference plasmid before fold normalization. Bars show means ± SD (WT, n = 4; each mutant, n = 3 per treatment). One-sided, unpaired Student’s *t* tests assuming equal variances were used for C; the same tests with Holm correction within each graph were used for D–F. Comparisons were with WT in C and E and between 0 and 2 mM Put within each construct in D and F. *, *P* < 0.05; **, *P* < 0.01; ***, *P* < 0.001; n.s., *P* ≥ 0.05.

To test whether translation of these uORFs is required for Put-responsive regulation, the start codons of these uORFs in the *ADC2* 5’-UTR:*Eluc* reporter were replaced by codons that were not expected to function as start codons (AUG to AAA, AUU to GUU, and CUG to CUC) (Fig. 4B). Our previous study using Arabidopsis suspension cells suggested that both the AUU and CUG codons of the non-AUG-initiated uORF contribute to repression of the main ORF (Hiragori et al. 2023). We therefore mutated either or both of these codons. WT and mutant reporter constructs were introduced into Arabidopsis mesophyll protoplasts, and Eluc activity was compared in the presence and absence of exogenous Put. Mutation of the AUG codon increased Eluc activity in the absence of exogenous Put, indicating that the AUG-initiated uORF represses expression of the main ORF (Fig. 4C). However, this mutation did not affect the Put-responsive reduction in Eluc activity (Fig. 4D). Thus, the AUG-initiated uORF contributes to basal repression of the main ORF but is not required for Put-responsive regulation.

Under untreated conditions, mutation of the CUG codon or simultaneous mutation of the AUU and CUG codons increased Eluc activity relative to the WT reporter, whereas mutation of the AUU codon alone had no significant effect on Eluc activity (Fig. 4E). By contrast, none of these non-AUG-codon mutants showed a significant reduction in Eluc activity following Put treatment (Fig. 4F). Thus, disruption of either the AUU or CUG initiation codon abolished the Put-responsive reduction in Eluc activity. Because the AUU and CUG codons are located close to the ADC-box, we assessed whether these mutations altered the predicted secondary structure of the ADC-box region using an *in silico* RNA secondary structure prediction tool (Sato et al. 2021). The single-codon mutations of either AUU or CUG had little effect on the predicted RNA secondary structure, whereas simultaneous mutation of both codons altered the predicted structure (Supplemental Fig. S3). Although the effects of the substitutions on RNA structure cannot be completely excluded, the loss of Put-responsive regulation in each of the single mutants, whose predicted structures remained largely unchanged, supports a requirement for both initiation codons in the response under these assay conditions, consistent with a role for uORF translation.

### Predicted RNA secondary structure in the ADC-box region contributes to Put-responsive repression

We next tested whether the ADC-box is also required for Put-responsive regulation mediated by the *ADC2* 5’-UTR. To distinguish the contribution of its conserved nucleotide sequence from that of its predicted RNA secondary structure, we generated an “unpaired” mutant designed to disrupt base pairing within the ADC-box and a “paired” mutant carrying additional nucleotide substitutions designed to restore the predicted secondary structure (Fig. 5, A and B). Although the unpaired and paired mutations altered part of the amino acid sequence of the overlapping AUU/CUG-initiated uORF, neither mutation changed the position or length of the uORF. Under untreated conditions, the unpaired and paired mutants showed approximately eightfold and fourfold the Eluc activity of the WT reporter, respectively (Fig. 5C). However, the unpaired mutant showed no significant Put-dependent decrease in Eluc activity, whereas the paired mutant retained Put-responsive repression comparable to that of the WT reporter, with activity decreasing to approximately 25% of the untreated level (Fig. 5D). The paired mutant therefore restored Put responsiveness despite extensive changes in its primary sequence. Together, these results indicate that the RNA secondary structure formed in the ADC-box region is critical for Put-responsive regulation.

**Figure 5.**
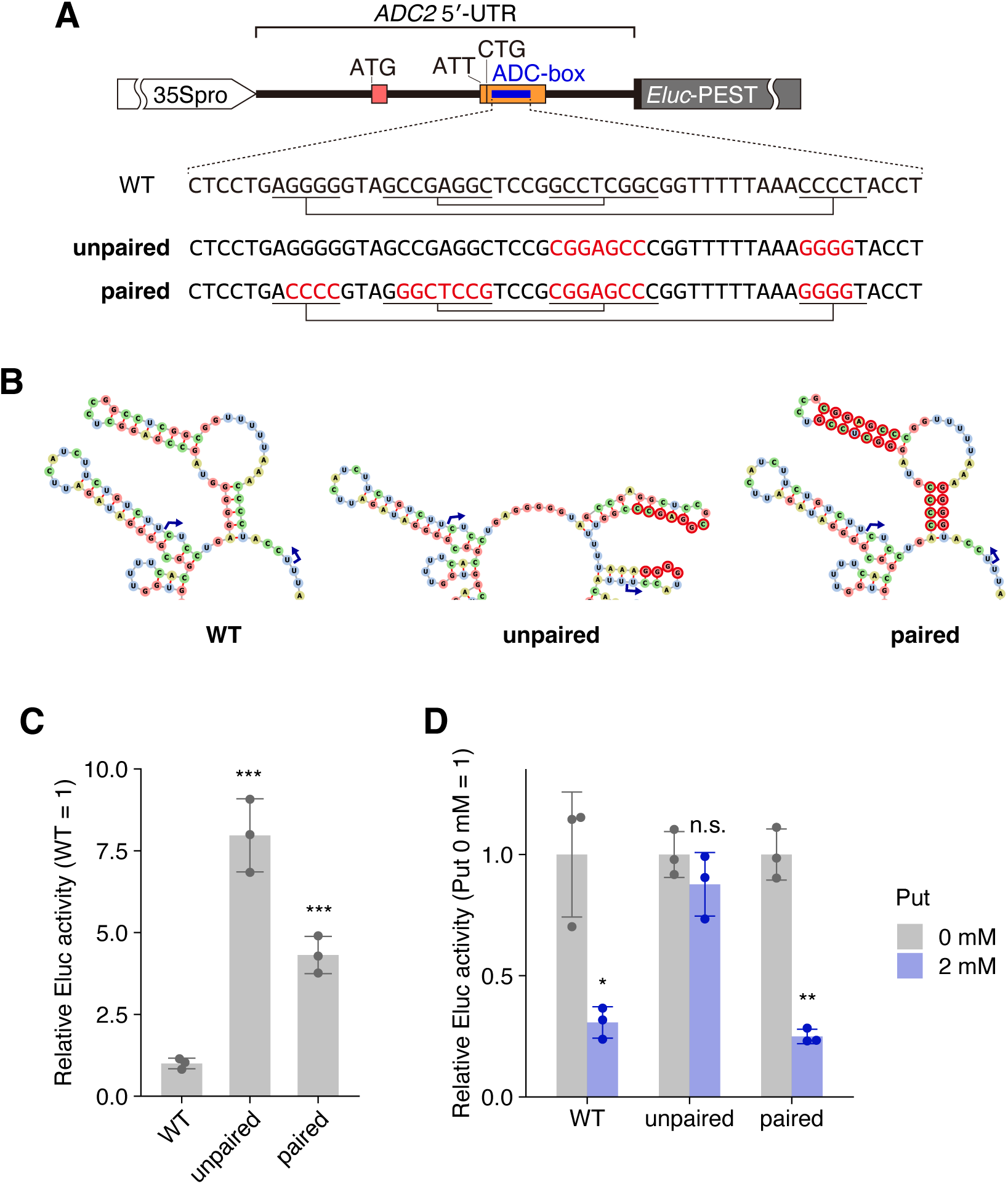
Effects of disrupting and restoring the predicted RNA structure in the ADC-box region on Put-responsive repression. (A) Reporter design and WT, unpaired, and paired ADC-box sequences. Substituted nucleotides are shown in red, and connecting lines indicate predicted base-paired segments. The mutations overlap the non-AUG-initiated uORF. (B) RNA secondary structures predicted using MXFold2. Blue arrows mark the boundaries of the ADC-box; base colors distinguish nucleotide identities, and red outlines mark substituted nucleotides. (C) Relative Eluc activity under untreated conditions, with WT set to 1. (D) Relative Eluc activity after treatment with 0 or 2 mM Put, with the untreated condition set to 1 for each construct. Reporters were tested in mesophyll protoplasts incubated at 22°C for 24 h in the dark. Eluc activity was normalized to Rluc activity from the reference plasmid before scaling to the control-group mean within each experiment. Bars show means ± SD (n = 3 per group). One-sided, unpaired Student’s *t* tests assuming equal variances were used with Holm correction within each graph: each mutant was compared with WT in C, and treated and untreated samples were compared within each construct in D. *, *P* < 0.05; **, *P* < 0.01; ***, *P* < 0.001; n.s., *P* ≥ 0.05.

### Regulation mediated by the *ADC2* 5’-UTR contributes to normal growth and Put homeostasis in Arabidopsis

The expression level of *ADC2* is a major determinant of Put levels in Arabidopsis (Urano et al. 2004; Rossi et al. 2015). Put-responsive repression of *ADC2* expression may therefore function as a negative feedback mechanism that contributes to Put homeostasis. To examine the importance of *ADC2* 5’-UTR-mediated regulation *in planta*, we introduced mutations into the genomic region corresponding to the *ADC2* 5’-UTR using the CRISPR-Cas9 system. We obtained two mutant lines, #3-7 and #11-8, harboring 25-and 31-nt deletions, respectively (Fig. 6A; Supplemental Fig. S4A). In both mutants, the deleted regions span the AUU/CUG-initiated uORF and the ADC-box. Both deletions shifted the reading frame of the AUU/CUG-initiated uORF, changing the position of the stop codon 17 nt downstream of the original stop codon (Fig. 6A; Supplemental Fig. S4A). Because the deleted nucleotides participate in base pairing within the ADC-box RNA secondary structure, we predicted the RNA secondary structures surrounding the ADC-box in each mutant sequence *in silico*. The predicted structures of both mutant sequences differed substantially from that of the WT sequence (Supplemental Fig. S4B). When grown on half-strength Murashige and Skoog (MS) medium, #3-7 and #11-8 showed significantly reduced shoot fresh weight and primary root length compared with Col-0, with #11-8 exhibiting the more severe growth phenotype (Fig. 6, B–D). Radicle emergence and cotyledon expansion were delayed in both lines, and the delay in cotyledon expansion was particularly pronounced in #11-8 (Supplemental Fig. S5, A and B).

**Figure 6.**
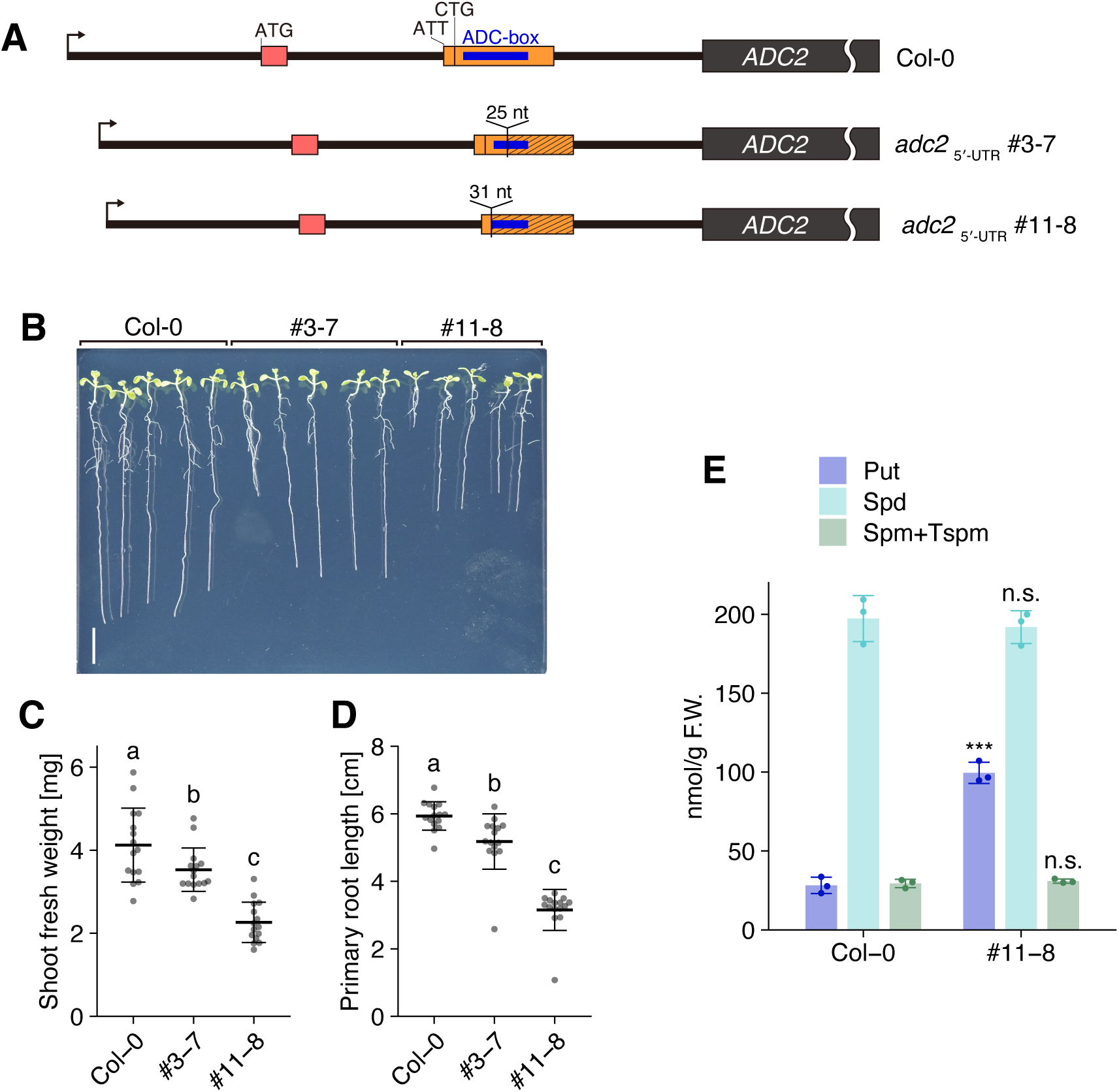
*ADC2* 5’-UTR genome-edited lines exhibit growth defects, and #11-8 accumulates Put. (A) Organization of the WT and edited *ADC2* 5’-UTRs. Lines #3-7 and #11-8 contain 25- and 31-nt deletions, respectively, overlapping with the non-AUG-initiated uORF and ADC-box. Codons are shown in DNA notation. (B) Seedlings of Col-0, #3-7, and #11-8 grown vertically on half-strength MS medium solidified with gellan gum, photographed on day 10 after stratification. Scale bar, 1 cm. (C) Shoot fresh weight and (D) primary root length measured on day 10. Seedlings that had germinated by day 2 were transferred on day 5 to three fresh plates, with five seedlings per genotype per plate (n = 15 per genotype). Points represent individual seedlings; thick horizontal lines indicate means and thin lines indicate SD ranges. Different letters indicate significant differences among genotypes (one-way ANOVA followed by Tukey’s test, *P* < 0.05). (E) Put, Spd, and combined Spm+Tspm abundance in Col-0 and #11-8, determined by HPLC in 14-day-old whole seedlings grown on half-strength MS agar medium and expressed per gram fresh weight (F.W.). The combined Spm+Tspm peak was quantified using the Spm calibration curve. Bars show means ± SD of three samples per genotype, each comprising approximately 20 seedlings pooled from one plate (approximately 100 mg). Comparisons with Col-0 used one-sided, unpaired Student’s *t* tests assuming equal variances, with Holm correction across the comparisons in E. *, *P* < 0.05; **, *P* < 0.01; ***, *P* < 0.001; n.s., *P* ≥ 0.05.

We quantified polyamines in Col-0 and #11-8 by high-performance liquid chromatography (HPLC). In #11-8, Put abundance was approximately 3.5 times that in Col-0, whereas Spd and the combined Spm+Tspm fraction showed no significant differences (Fig. 6E; Supplemental Fig. S6). Put accumulation in #11-8, the line with the more severe growth and germination phenotypes, suggests an association between Put accumulation and these developmental defects. Together, these results support a contribution of *ADC2* 5’-UTR-mediated regulation to Put homeostasis and normal germination and growth in Arabidopsis.

## DISCUSSION

### Put-responsive feedback regulation mediated by the *ADC2* 5’-UTR

Our findings identify the Arabidopsis *ADC2* 5’-UTR as a *cis*-regulatory module that mediates negative feedback regulation in response to Put availability. Exogenous Put reduced Eluc activity in mesophyll protoplasts, whereas *adc2* mutations or pharmacological inhibition of ADC increased it; Put supplementation counteracted these increases (Fig. 2). In transgenic plants, Put reduced Eluc activity more strongly than reporter mRNA abundance and decreased calculated relative Eluc TE, supporting a major contribution from translational repression (Fig. 3).

Exogenous Put reduced the abundance of reporter mRNA modestly but significantly in an *ADC2* 5’-UTR-dependent manner (Fig. 3C). Because Put did not affect the abundance of *MASP1* reporter mRNA driven by the same 35Spro, this reduction may reflect *ADC2* 5’-UTR-dependent mRNA destabilization rather than a change in transcription. Translation termination at a uORF or ribosome stalling can engage mRNA quality control pathways, including NMD (Gaba et al. 2005; Nyikó et al. 2009; Uchiyama-Kadokura et al. 2014). Changes in ribosome behavior on the *ADC2* 5’-UTR could therefore couple repression of main ORF translation to reduced mRNA stability.

### Roles of the *cis*-regulatory elements in the *ADC2* 5’-UTR and potential mechanisms of Put-responsive regulation

Mutational analyses showed that the AUU/CUG-initiated uORF and the predicted RNA secondary structure of the ADC-box region are central to Put-responsive regulation, whereas the short AUG-initiated uORF is dispensable for this response (Figs. 4 and 5). Elimination of either the AUU or CUG initiation codon abolished the Put-responsive repression, as did disruption of predicted base pairing in the ADC-box region (Fig. 4F; Fig. 5D). Conversely, a compensatory mutation predicted to reconstitute the ADC-box structure reinstated the Put-responsive repression despite extensive changes in primary sequence (Fig. 5), supporting the importance of the RNA structure rather than the conserved sequence itself. Notably, the effects of these mutations on Put responsiveness did not parallel their effects on basal repression. Mutation of the AUG-initiated uORF increased basal Eluc activity without abolishing Put-responsive repression; mutations of the AUU/CUG-initiated uORF abolished Put responsiveness despite differing effects on basal activity; and both the unpaired and paired ADC-box mutants increased basal activity, although only the unpaired mutant lost Put responsiveness (Figs. 4 and 5). Together, these findings indicate that Put-responsive regulation does not simply reflect the strengthening or alleviation of basal repression.

Because the AUU/CUG-initiated uORF overlaps the ADC-box (Fig. 4A; Supplemental Fig. S2), their contributions to Put responsiveness cannot be completely separated. Nevertheless, the paired mutant restored Put responsiveness despite containing more extensive sequence changes within the overlapping uORF than the unpaired mutant, arguing that the loss of responsiveness in the unpaired mutant cannot be explained simply by changes in the uORF sequence (Fig. 5). Conversely, the ΔAUU and ΔCUG mutations abolished Put responsiveness, although both mutations were predicted to have little effect on the ADC-box RNA structure, making it unlikely that their effects resulted from disruption of this structure (Fig. 4F; Supplemental Fig. S3). These observations support a model in which both the AUU/CUG-initiated uORF and the ADC-box RNA structure are required for Put-responsive repression.

One possible mechanism underlying this Put-responsive regulation is that Put stabilizes an RNA structure in the ADC-box region. Owing to their cationic nature, polyamines can stabilize RNA structures by interacting with the negatively charged phosphate backbone (Kusano et al. 2008; Igarashi and Kashiwagi 2010). A stabilized ADC-box structure could repress main ORF translation in at least two ways. First, because it lies near and downstream of the AUU/CUG initiation sites, it could slow scanning by a 43S preinitiation complex and thereby enhance recognition of either of the non-AUG start codons, consistent with previous reports that downstream RNA structures can promote initiation at upstream near-cognate codons (Kozak 1990; Wang et al. 2022). Increased uORF initiation would reduce the number of ribosomes reaching the main ORF. Another possibility is that the ADC-box structure could slow elongation or termination of ribosomes translating the uORF, obstructing subsequent scanning or translating ribosomes and reducing the number of ribosomes that reach the main ORF (Dever et al. 2020; Bao et al. 2022). In either model, cooperation between the ADC-box RNA structure and the AUU/CUG-initiated uORF could explain the effects of the mutations on Put responsiveness (Figs. 4 and 5), although the current data do not distinguish between these models.

Put could also act directly on ribosomes to selectively modulate *ADC2* translation through its 5’-UTR. Put has been reported to associate with actively translating ribosomes and to alter their ribosomal protein composition (Murillo et al. 2025). Tspm can also bind to the peptidyl transferase center in ribosomes and modulate translation of specific genes (Ko et al. 2026). These findings leave open the possibility that direct interaction between Put and ribosomes modulates translation on the *ADC2* 5’-UTR, although direct evidence for such a mechanism is currently lacking.

### Physiological significance of regulation mediated by the *ADC2* 5’-UTR

The genome-edited lines #3-7 and #11-8 provide a physiological link between regulation mediated by the *ADC2* 5’-UTR and Put homeostasis. Both lines exhibited impaired growth and delayed germination, with more pronounced defects in #11-8 (Fig. 6; Supplemental Fig. S5). The increased Put level in this more severely affected line suggests that Put overaccumulation may be associated with the developmental defects (Fig. 6E).

The phenotypes of the *ADC2* 5’-UTR mutants are consistent with a previous report showing that plants constitutively overexpressing *ADC2* overaccumulate Put and show growth defects associated with reduced levels of bioactive gibberellins and their precursors (Alcázar et al. 2005). Because gibberellins are phytohormones that regulate the timing of seed germination, the delayed germination of the 5’-UTR mutants could involve altered gibberellin metabolism. Together, these findings support a contribution of the regulatory functions of the *ADC2* 5’-UTR to Put homeostasis and normal germination and growth *in planta*.

The absence of corresponding increases in the other major polyamines (Fig. 6E) suggests that regulation mediated by the *ADC2* 5’-UTR is particularly important for restricting Put accumulation rather than uniformly controlling downstream polyamine levels. Together with feedback mechanisms regulating *SAMDC* and *ACL5* expression, this finding supports a model in which polyamine homeostasis is controlled separately at multiple steps of the biosynthetic pathway.

### Potential conservation and lineage-dependent importance of the *ADC* 5’-UTR-mediated regulation

Conservation of the ADC-box and the non-AUG-initiated uORF in multiple plant *ADC* genes raises the possibility that feedback regulation mediated by *ADC* 5’-UTRs contributes to Put homeostasis beyond Arabidopsis (Wu et al. 2019; Hiragori et al. 2023). However, the physiological contribution of this feedback regulation may depend on the organization of Put biosynthesis in each lineage. Arabidopsis lacks a functional ODC, and *ADC2* is a major determinant of its Put levels (Fig. 1; Hanfrey et al. 2001; Urano et al. 2004; Rossi et al. 2015), potentially making feedback through the *ADC2* 5’-UTR particularly consequential. In plants possessing both ADC and ODC pathways, regulation of *ODC* genes could complement or partly substitute for regulation mediated by *ADC* 5’-UTRs. A more complete understanding of Put homeostasis in plants will require comparative analyses of the regulation of genes involved in Put biosynthesis and metabolism across species with different biosynthetic architectures.

## MATERIALS AND METHODS

### Plant materials and growth conditions

*Arabidopsis thaliana* Col-0 was used as WT. The T-DNA insertion lines *adc2-1* (SALK_073977) and *adc2-2* (WiscDsLoxHs065_09H), both in the Col-0 background, were obtained from the Arabidopsis Biological Resource Center (Columbus, OH, USA). Homozygous T-DNA insertions were confirmed by the absence of a gene-specific PCR product and the presence of a gene-specific/T-DNA-border PCR product. Primers used for genotyping are listed in Supplemental Table S1. MM2d suspension cells were derived from Landsberg *erecta*. Stable lines carrying an *ADC2* 5’-UTR or *MASP1* 5’-UTR reporter and two genome-edited lines #3-7 and #11-8, carrying 25- and 31-nt deletions in the *ADC2* 5’-UTR, respectively, were generated in this study (see below).

Seeds were surface-sterilized with sodium hypochlorite and stratified at 4°C in the dark for 4–6 d for transgenic reporter assays or 6 d for germination and growth assays. Seedlings were grown at 22°C under continuous LED light (75–1OO µmol m^−^² s^−^¹) on half-strength MS medium containing 1% (w/v) sucrose and O.5 g L^−^¹ 2-(N-morpholino)ethanesulfonic acid (MES; pH 5.7), solidified with 0.8% (w/v) agar or, for growth measurements, 0.5% (w/v) gellan gum. Mesophyll protoplast donors were grown for 4 weeks in Jiffy-7 pellets (Sakata Seed Corporation, Yokohama, Japan) at 22°C under continuous fluorescent light (50–8O µmol m^−^² s^−^¹).

### Plasmid construction

Plasmids used or generated in this study and primers used for plasmid construction are listed in Supplemental Tables S2–S4. Reporter plasmids used for transient expression assays were derived from pKM56, which contains a 35Spro, an *Eluc*-PEST, and the Arabidopsis *HSP18.2* terminator (*HSP*ter) in pUC19 (Hayashi et al. 2017; Hiragori et al. 2023). pKM56 served as the control plasmid. For the *ADC2* 5’-UTR:*Eluc* reporter plasmid pYH21, the *ADC2* 5’-UTR and first 5 nt of the main ORF were amplified from Col-0 cDNA and inserted between the XbaI and SalI sites of pKM56 by seamless ligation cloning extract (SLiCE) assembly (Motohashi 2017). Mutations were introduced by overlap PCR (Ho et al. 1989). All PCR-amplified regions were verified by Sanger sequencing. The reference plasmid pKN17, which carries *Rluc* under the control of 35Spro, was generated by inserting annealed complementary oligonucleotides “RLUC 5UTR Fw/Rv” between the XbaI and SalI sites of pKM5 (Hayashi et al. 2017) and was used in all transient assays.

The binary vector pYH74 was derived from pKI1.1R (Tsutsui and Higashiyama 2017). An annealed “GG to CC IF for/rev” fragment was inserted into SbfI/PmeI-digested pKI1.1R by In-Fusion assembly (Takara, Shiga, Japan) to remove SbfI and SalI sites. The resulting plasmid was digested with KpnI and StuI. A fragment amplified from pKI1.1R by sequential PCRs with “CBSAE_for2_new”/“CBSAE_rev” and “CBSAE_for”/”CBSAE_rev” was inserted into the KpnI/StuI sites by In-Fusion assembly, replacing the Cas9/guide RNA (gRNA) cassette with an SbfI–BamHI–EcoRI–SmaI–XmaI multiple cloning site (MCS). To generate the *MASP1* 5’-UTR reporter plasmid pYH76, the SbfI/EcoRI fragment containing the 35Spro::*MASP1* 5’-UTR:*Eluc*-PEST::*HSP*ter cassette was excised from pYH17 (Hiragori et al. 2023) and ligated into SbfI/EcoRI-digested pYH74. To create the *ADC2* 5’-UTR reporter plasmid pYH75, the SbfI/BglII fragment containing the *ADC2* 5’-UTR:*Eluc* construct was excised from pYH21 and ligated into SbfI/BglII-digested pYH76.

### Generation of transgenic and genome-edited lines

Constructs were introduced into *A. thaliana* plants by the floral dip method (Clough and Bent 1998) using *Agrobacterium tumefaciens* strain GV2260 (Deblaere et al. 1985). For transgenic reporter lines, hygromycin B-resistant T1 plants were selected. Then, T2 lines carrying the transgene at a single locus were selected based on a 3:1 segregation ratio of hygromycin B resistance, and homozygous lines were established in the T3 generation by confirming that no segregation occurred. Two *ADC2* 5’-UTR reporter lines #1 and #2 originated from different T0 plants.

For genome-edited lines, annealed “SPE2 gR1 F/R” or “SPE2 gR2 F/R” oligonucleotides (Supplemental Table S5) were ligated into AarI-digested pKI1.1R to generate pYH70 (gRNA1) or pYH71 (gRNA2). T1 transformants were selected by hygromycin B resistance. CRISPR/Cas9-induced mutations were detected by heteroduplex mobility assays (Foster et al. 2019) in T1 and T2 plants and verified by Sanger sequencing. Homozygous T3 plants or their progeny were used for analyses. Homozygous mutant lines #3-7 and #11-8 were generated using gRNA2 and gRNA1, respectively.

### Mesophyll protoplast preparation and transfection

Mesophyll protoplasts were isolated following Wu et al. (2009), with some modifications. Lower epidermal layers of rosette leaves were removed using masking tape (Kamoi Kakoshi, Okayama, Japan). Twenty leaves were incubated in 20 mL of enzyme solution (1% Cellulase “Onozuka” R10 and 0.25% Macerozyme R10 (Yakult, Tokyo, Japan), 0.4 M mannitol, 10 mM CaCl2, 2O mM KCl, O.1% bovine serum albumin, and 20 mM MES-KOH, pH 5.6) for 2–3 h at room temperature and 40 rpm. Centrifugation during washing and after polyethylene glycol (PEG) treatment was performed at 1,000 rpm for 3 min at 4°C using a Sorvall ST 8FR centrifuge (Thermo Fisher Scientific, Waltham, MA, USA) with a swinging-bucket rotor. W5 buffer contained 154 mM NaCl, 125 mM CaCl2, 5 mM KCl, 5 mM glucose, and 2 mM MES (pH 5.7).

Each transfection contained 1.5 pmol each of the reporter and reference plasmids in 10 µL water, 1.5 × 1Oၫ protoplasts in 1OO µL MaMg solution (O.4 M mannitol, 15 mM MgCl2, and 4 mM MES-KOH, pH 5.6), and 11O µL PEG solution (4O% PEG4OOO, O.1 M CaCl2, and O.2 M mannitol). After 15 min at room temperature, 800 µL W5 was added. Protoplasts were pelleted and resuspended in 200 µL WI medium (0.5 M mannitol, 20 mM KCl, and 4 mM MES-KOH, pH 5.6). Put was added to the protoplast cultures at 0–4 mM for dose-response assays or at 0 or 2 mM for other comparisons. The protoplasts were then incubated at 22°C for 24 h in the dark.

### MM2d suspension cell protoplast preparation and transfection

MM2d protoplast isolation and transfection followed Hiragori et al. (2023). Briefly, cells harvested 3 d after subculture were digested in modified Linsmaier and Skoog (LS) medium containing 1% Cellulase “Onozuka” RS (Yakult), 0.1% Pectolyase Y23 (Kyowakasei, Osaka, Japan), and 0.4 M mannitol at 26°C. Each transfection contained 3 µg of pYH21 and pKN17 in 6 µL water, 3 × 1Oၫ protoplasts in 1OO µL MaMg solution (O.4 M mannitol, 15 mM MgCl2, and 5 mM MES, pH 5.8), and 106 µL PEG solution. After PEG treatment, cells were resuspended in 200 µL modified LS medium containing 0.4 M mannitol. Put and/or DFMA were added to final concentrations of 1 mM each immediately before incubation at 22°C for 24 h in the dark.

### Luciferase assay

For transient expression assays, protoplasts were frozen in liquid nitrogen and extracted by vortexing for 2 min in 100 or 200 µL buffer containing 100 mM sodium phosphate (pH 7) and 5 mM dithiothreitol (DTT). Eluc and Rluc activities were measured using the Dual-Luciferase Reporter Assay System (Promega, Madison, WI, USA), and Eluc activity was normalized to Rluc activity.

For transgenic reporter assays, 15 ten-day-old seedlings per flask were submerged in 10 mL liquid half-strength MS medium supplemented with 0 or 2 mM Put and shaken at 80 rpm for 24 h. Whole seedlings from each flask were pooled, ground in liquid nitrogen with a mortar and pestle, and divided for RNA and protein extraction. For protein extraction, the samples were vortexed for 2 min in the same extraction buffer. Eluc activity was measured using the Luciferase Assay System (Promega) and normalized to total protein determined with Pierce 660 nm Protein Assay Reagent (Thermo Fisher Scientific).

### RNA analysis by RT-qPCR

RNA was extracted using TRIzol (Thermo Fisher Scientific) and treated with RQ1 RNase-Free DNase (Promega). RNA (500 ng) was reverse-transcribed using SuperScript IV (Thermo Fisher Scientific) according to the manufacturer’s protocol, except that RNaseOUT was omitted, 100 ng of oligo(dT)ı2–ı8 primer was used, and reverse transcription was performed at 50°C for 60 min. RT-qPCR used LightCycler FastStart DNA MasterPLUS SYBR Green I and a LightCycler 2.0 instrument (Roche, Mannheim, Germany). *Eluc* transcript abundance was normalized to *EF1a* (At5g60390) in the same cDNA sample. For analysis of endogenous *ADC2* expression in Col-0 and the *adc2* mutants, three fresh rosette leaves from an untreated 4-week-old plant were pooled per biological replicate (n = 3 per genotype). *ADC2* transcript abundance was normalized to *EF1a*. Primers used for RT-qPCR are listed in Supplemental Table S6.

### RNA secondary structure prediction

WT and mutant RNA structures were predicted and drawn using the MXFold2 Server (https://ws.sato-lab.org/mxfold2/; Sato et al. 2021) with default settings. Inputs comprised the ADC-box region and 100 nt on each side.

### Growth and germination assays

For growth assays, seedlings that had germinated by day 2 after stratification were selected and transferred to fresh gellan-gum plates on day 5. On day 10, shoots were cut at the root–hypocotyl boundary and weighed individually in 1.5-mL tubes. Primary root length was measured from images using ImageJ.

For germination assays, seeds stored dry with silica gel in the dark for more than one month were sown on three agar plates. The same seeds were observed over 7 d after stratification. The percentage in each plate was treated as one replicate at each time point (n = 3).

### Polyamine quantification

Whole seedlings grown for 14 d on half-strength MS agar under continuous light (approximately 75 µmol m^−^² s^−^¹) were pooled by plate (approximately 2O seedlings, 1OO mg per sample; n = 3 per genotype). Frozen, ground tissue was extracted with four volumes (v/w) of 5% perchloric acid. A 200-µL supernatant aliquot was supplemented with 1,7-diaminoheptane (Tokyo Chemical Industry, Tokyo, Japan) to 50 µM in the extract, mixed with 400 µL of 10 mg mL^−^¹ dansyl chloride (Tokyo Chemical Industry) in acetone and 200 µL saturated sodium carbonate, and incubated overnight at room temperature in the dark. After adding 100 µL of 1OO mg mL^−^¹ aqueous proline and incubating for 3O min, samples were mixed with 25O µL benzene. The organic phase was recovered, dried under reduced pressure, dissolved in 100 µL acetonitrile:water (1:1), and filtered through a Millex syringe filter (0.45 µm; Merck Millipore, Burlington, MA, USA). Mixed Put, Spd, and Spm standards (each at 1, 10, 20, 40, or 80 µM in perchloric acid solution) were processed identically.

HPLC used a Puresil 5-µm C18 column (120 Å, 4.6 × 150 mm; Waters, Milford, MA, USA) with a Resolve Silica Guard-Pak precolumn (Waters), an ELITE LaChrom L-2130 pump and L-2400 UV detector, and D-2000 Elite software (Hitachi, Tokyo, Japan). Injection volume was 4O µL, detection was at 34O nm, and flow rate was 1 mL min^−^¹. Acetonitrile:water was 60:40 for 0–15 min, increased to 100% acetonitrile over 15–30 min, maintained for 30–45 min, and returned to 60:40 for 45–60 min.

### Sequence identifiers

The Arabidopsis locus identifiers discussed in this study are At4g34710 (*ADC2*), At2g16500 (*ADC1*), At4g03260 (*MASP1*), and At5g60390 (*EF1a*).

## SUPPLEMENTARY MATERIAL

Supplemental Figure S1. T-DNA insertion alleles of *ADC2*.

Supplemental Figure S2. Sequence and regulatory elements of the *ADC2* 5’-UTR.

Supplemental Figure S3. Predicted RNA secondary structures of non-AUG start-codon mutants.

Supplemental Figure S4. Deletion sequences and predicted RNA structures of *ADC2* 5’-UTR genome-edited alleles.

Supplemental Figure S5. Germination and cotyledon expansion of *ADC2* 5’-UTR genome-edited lines.

Supplemental Figure S6. HPLC chromatograms and calibration curves for polyamine quantification.

Supplemental Table S1. Primers used for mutant genotyping.

Supplemental Table S2. Plasmids used in transient assays and primers used for plasmid construction.

Supplemental Table S3. Plasmids used for generation of transgenic or genome-edited lines and oligo DNA used for plasmid construction.

Supplemental Table S4. Primers used for plasmid construction. Supplemental Table S5. Oligo DNA used for guide RNA cloning. Supplemental Table S6. Primers used for RT-qPCR.

## ACKNOWLEDGMENTS

The authors thank Ms. Saeko Yasokawa for technical assistance, and Ms. Sayo Aono and Ms. Aki Kosugi for general assistance.

## AUTHOR CONTRIBUTIONS

Y.H. and H.O. designed research. Y.H., M.Y., T.K., A.K., Y.S., and Y.G. performed the experiments. Y.H., Y.S., Y.Y., S.N., and H.O. analyzed data. Y.H., S.N., and H.O. wrote the manuscript.

## Abbreviations

5’-UTR, 5’ untranslated region, 35Spro, cauliflower mosaic virus 35S RNA promoter, ACL5, ACAULIS5, ADC, arginine decarboxylase, ADC1, arginine decarboxylase 1, ADC2, arginine decarboxylase 2, Arg, L-arginine, Col-0, Columbia-0, DFMA, a-difluoromethylarginine, Eluc, emerald luciferase, gRNA, guide RNA, HPLC, high-performance liquid chromatography, NMD, nonsense-mediated decay, ODC, ornithine decarboxylase, Orn, L-ornithine, Put, putrescine, Rluc, Renilla luciferase, RT-qPCR, reverse transcription quantitative PCR, SAMDC, *S*-adenosylmethionine decarboxylase, Spd, spermidine, Spm, spermine, TE, translation efficiency, Tspm, thermospermine, uORF, upstream open reading frame, WT, wild type.

---

**Supplemental Figure S1. T-DNA insertion alleles of *ADC2*.** (A) Locations of the T-DNA insertions in *adc2-1* (SALK_073977) and *adc2-2* (WiscDsLoxHs065_09H). Black arrowheads indicate the positions and orientations of the primers used for RT-qPCR. (B) Relative *ADC2* mRNA levels measured by RT-qPCR in the indicated genotypes. Each biological replicate comprised fresh rosette leaves from a 4-week-old plant. *ADC2* transcript abundance was normalized to *EF1a* and expressed relative to Col-0. The bar and error bars for Col-0 represent the mean ± SD (n = 3). In *adc2-1* (†), only one of the three biological replicates yielded a quantifiable value, shown as a single dot, whereas the remaining two showed target-specific amplification below the lowest point of the standard curve. In *adc2-2* (‡), two replicates showed target-specific amplification below the lowest point of the standard curve, while only non-specific amplification was detected in the remaining replicate. Thus, no quantifiable values were obtained for *adc2-2*.

**Supplemental Figure S2. Sequence and regulatory elements of the *ADC2* 5’-UTR.** The nucleotide sequence is shown in DNA notation, with positions numbered from the first nucleotide. The first nucleotide shown in red marks the transcription start site (TSS) peak annotated in Plant Promoter Database 3.0 (http://ppdb.agr.gifu-u.ac.jp; Hieno et al. 2014). The short AUG-initiated uORF is highlighted in red and the AUU/CUG-initiated uORF in orange. The ADC-box is underlined in blue. Boxes mark the uORF initiation codons; the beginning of the main ORF is highlighted in dark gray.

**Supplemental Figure S3. Predicted RNA secondary structures of non-AUG start-codon mutants.** Predicted structures are shown for WT, ΔAUU, ΔCUG, and ΔAUUΔCUG sequences encompassing the ADC-box region and 100 nt on each side, calculated and drawn using MXFold2 Server with default settings. Blue arrows mark the ADC-box boundaries, base colors distinguish nucleotide identities, and red outlines mark substituted nucleotides.

**Supplemental Figure S4. Deletion sequences and predicted RNA structures of *ADC2* 5’-UTR genome-edited alleles.** (A) Alignment of the WT region with the 25-nt deletion in #3-7 and the 31-nt deletion in #11-8. Orange shading marks the uORFs; the frameshifted mutant uORFs terminate 17 nt downstream of the WT stop position. Black and gray boxes indicate guide RNA target regions and protospacer adjacent motifs (PAMs), respectively. Blue letters mark the ADC-box, red dashed segments indicate deleted nucleotides, and black underlines and connecting lines indicate predicted base pairing. (B) Structures predicted using MXFold2 for the WT and edited ADC-box regions with 100 nt on each side. Blue arrows mark the ADC-box boundaries, and base colors distinguish nucleotide identities. Red triangles mark deletion sites, with the numbers indicating deleted lengths.

**Supplemental Figure S5. Germination and cotyledon expansion of *ADC2* 5’-UTR genome-edited lines.** Germination scored by radicle emergence (A) and cotyledon expansion (B) in Col-0, #3-7, and #11-8 over 7 d after 6 d of stratification. Seeds were grown on half-strength MS agar medium at 22°C under continuous LED illumination (75–1OO µmol m^−^² s^−^¹). Germination was scored when the seed coat had split and the radicle protruded; cotyledon expansion was scored when the cotyledons were fully exposed and separated. Percentages were calculated using all seeds sown in one plate as the denominator. Each plate contained 76 seeds per genotype, and each plate-level proportion was one replicate (n = 3 plates). Values are means ± SD.

**Supplemental Figure S6. HPLC chromatograms and calibration curves for polyamine quantification.** Chromatograms are shown for mixed Put, Spd, and Spm standards and for three samples each of Col-0 and #11-8, labeled rep1–rep3, corresponding to the analysis in Fig. 6E. Each sample comprised approximately 20 whole seedlings pooled from one plate at 14 d of growth. Dansylated polyamines were detected by UV absorbance at 340 nm. Peaks are designated 0, 1,7-diaminoheptane (internal standard); 1, Put; 2, Spd; and 3, Spm in the standards and the unresolved Spm+Tspm fraction in plant samples. Relative peak area is the polyamine peak area divided by the internal-standard peak area. Calibration plots show relative peak area against concentration for the five nonzero standard concentrations, with fitted equations and coefficients of determination displayed in the figure. All sample relative peak areas were within the calibration ranges. Std, standard.

**Supplemental Table S1. Primers used for mutant genotyping.** Primer names, targets, and sequences are listed for the *ADC2* genome-edited region and the two T-DNA insertion lines. The gene-specific pairs SALK_073977 F/R and WiscDsLoxHs065_09H F/R were used together with gene-specific/T-DNA border primer combinations; LBb1.3 and LB4 modified were used for SALK_073977 and WiscDsLoxHs065_09H, respectively.

**Supplemental Table S2. Plasmids used in transient assays and primers used for plasmid construction.** Plasmid identifiers, reporter designs, and cloning primer names are listed. “Original plasmid” identifies the vector backbone used for insertion. All listed plasmids are based on pUC19. Primer sequences are provided in Supplemental Table S4.

Supplemental Table S3. Plasmids used for generation of transgenic or genome-edited lines and oligo DNA used for plasmid construction. *ADC2* and *MASP1* reporter constructs for transformation and CRISPR-Cas9 guide constructs for genome editing are listed with their vector and oligonucleotide identifiers. Guide RNA cloning oligonucleotide sequences are provided in Supplemental Table S5.

**Supplemental Table S4. Primers used for plasmid construction.** Primer sequences are given in the 5’-to-3’ direction. Primer names containing “SPE2” are retained to match the original laboratory records.

**Supplemental Table S5. Oligo DNA used for guide RNA cloning.** Sequences of SPE2 gR1 F/R and SPE2 gR2 F/R are listed in the 5’-to-3’ direction. Each oligonucleotide pair was annealed and ligated into *Aar*I-digested pKI1.1R to generate pYH70 (gRNA1) or pYH71 (gRNA2), respectively (Supplemental Table S3).

**Supplemental Table S6. Primers used for RT-qPCR.** Primer names, target genes, and sequences are listed for *Eluc*, *ADC2*, and *EF1a* (At5g60390). *EF1a* was used as the reference transcript for *Eluc* mRNA measurements in the stable reporter lines and for *ADC2* mRNA measurements in Supplemental Fig. S1B.

