## Supplemental Figure S1-6, Supplemental Table S1-6 for "Putrescine-responsive post-transcriptional feedback regulation of Arabidopsis *ADC2* mediated by 5’-UTR elements"

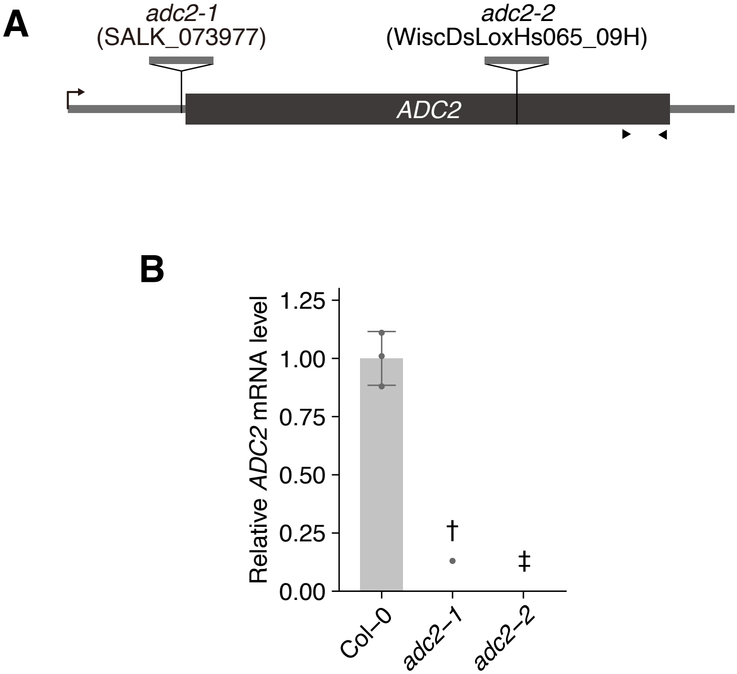


**Supplemental Figure S1**

1 ACGCCTTCTCTTCTTCTTCTTCTTCATTTCCGTCTTTTCCGCTAGGTTTCCGTCTCTGTTTTTTCTTGCATTTTC

76 CCCAGAATTCCTTCACATTCTCTCCTTTTCCTCTTTTTAATTCCTCACCACCTCAAATTCTTCATAAACAAATCC

159 179

151 ACTTCTGCATGGTTGGGAGAATCTGTTAGAGAATTTTTGTTGTTGTTGGGTTCAGATTTGGAAAATTTCCTTTTT

226 TTCGAGTTTCTTCTGAATTATTCATCGTGGAGATTCTCGGATCGGGAAGGGCTGCGAGAGGTGGTTTTCACGGCG

308 317 324

301 GGGATAGATTCATCTTCTGTCTTCTCCTGAGGGGGTAGCCGAGGCTCCGGCCTCGGCGGTTTTTAAACCCCTACC

376 397

376 TTTACAAATTCTGGGAAATTAGAAAAAAAAAGGTTTGATTTTTCAGTTTTTGCTCTGTTCTTCTCTCGAAAGTTT

520

451 TTGTTTTGTTCGAGAATTTCTGTAAAGTGAGATTGTTGATCGTTTGCTGATAAGAGGGTGAAGATAAAGATGCCT

**Supplemental Figure S2**

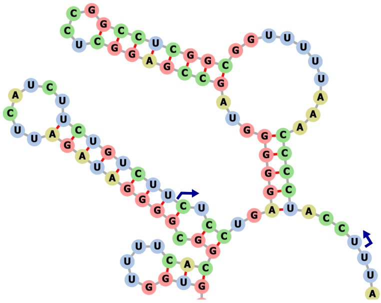

WT

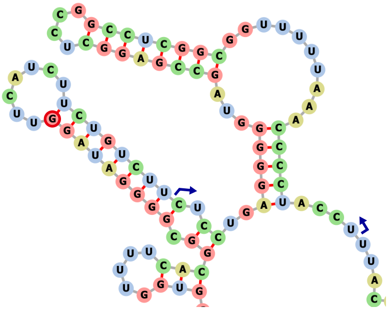

ΔAUU

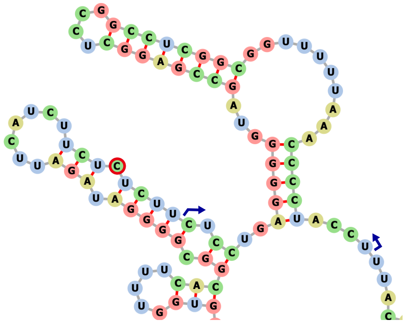

ΔCUG

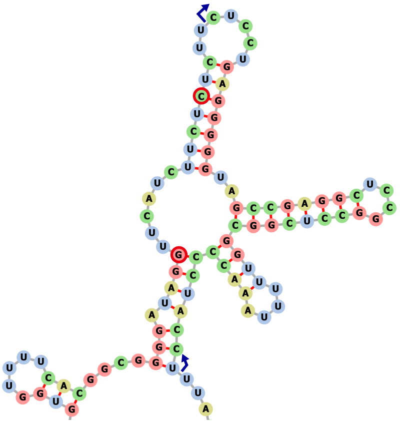

ΔAUUΔCUG

**Supplemental Figure S3**

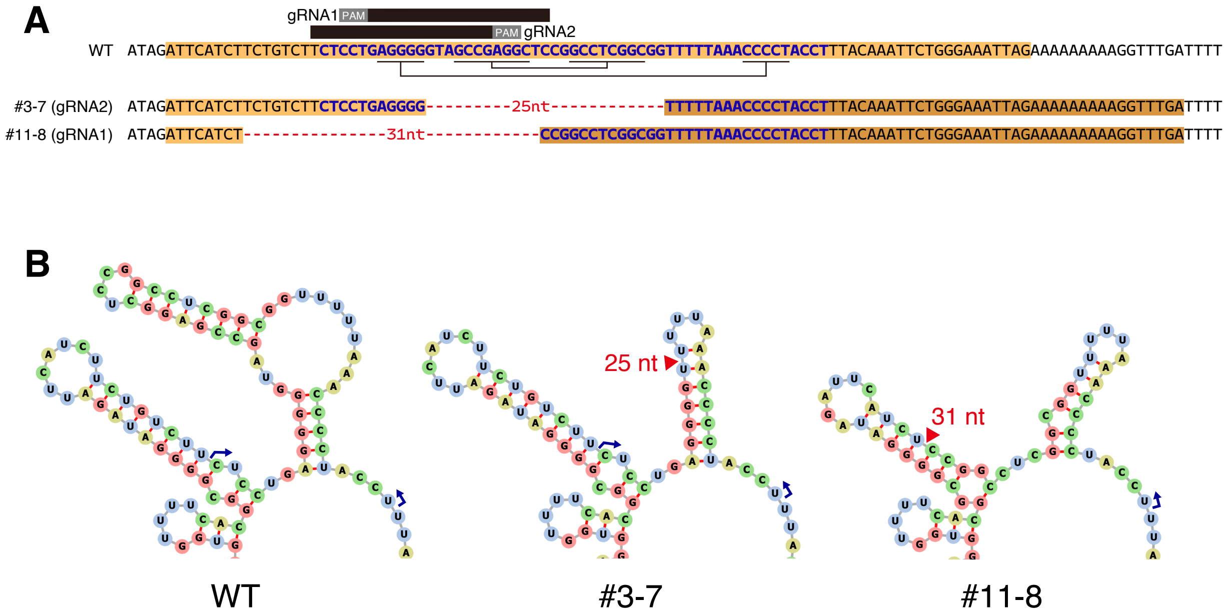


**Supplemental Figure S4**

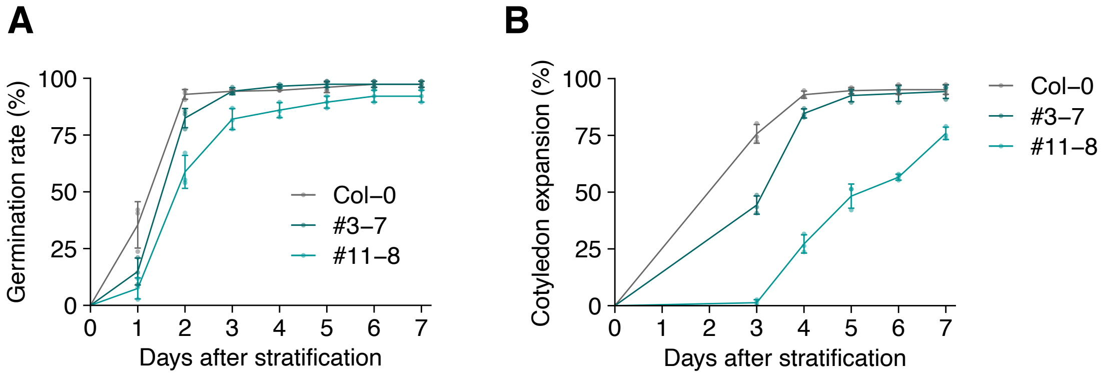


**Supplemental Figure S5**

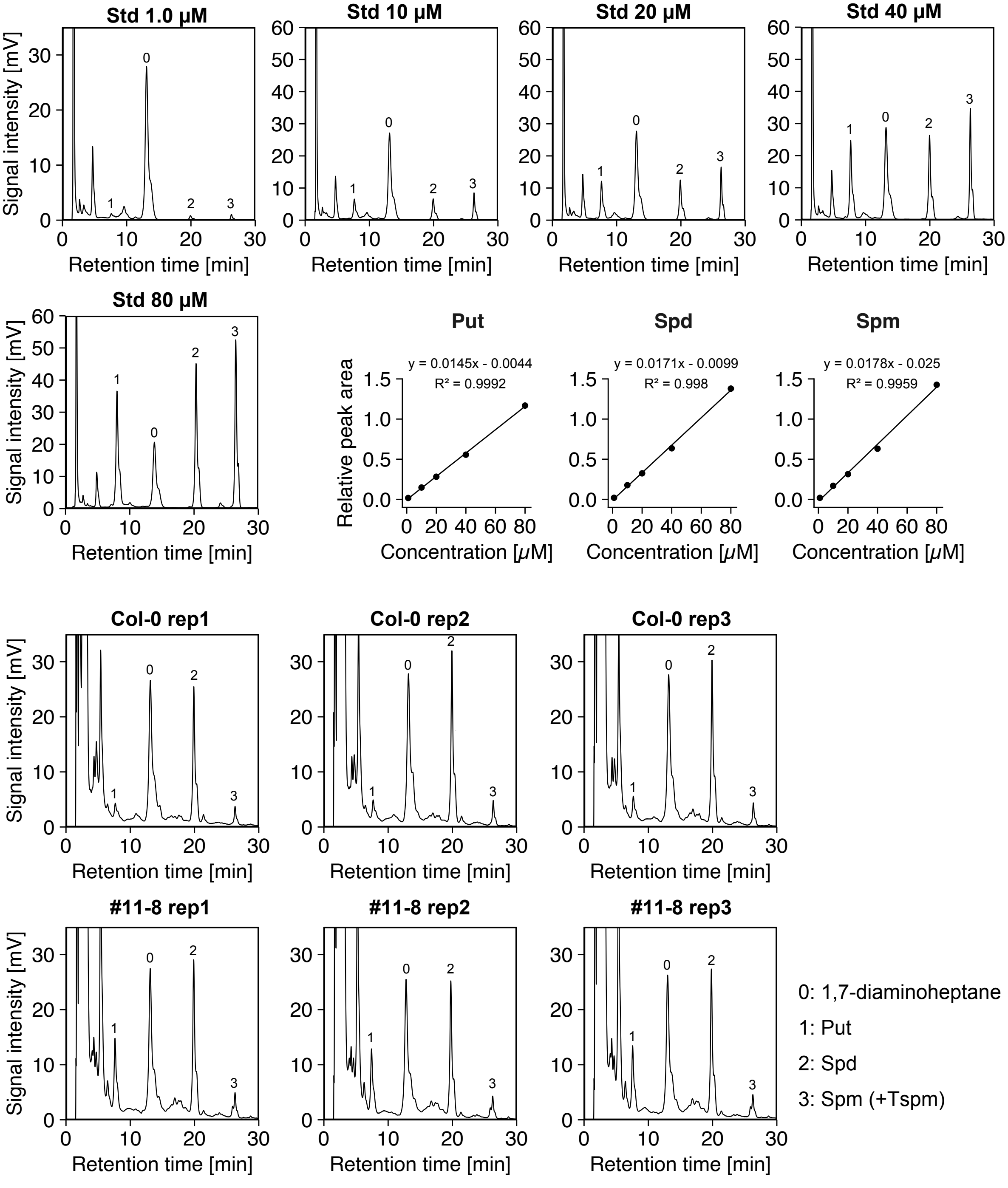


**Supplemental Figure S6**

**Supplemental Table S1. Primers used for mutant genotyping**

| Name | Target gene/transgene | Primer sequence (5′ to 3′) |
|---|---|---|
| SPE2 GE PCR F | *ADC2* | TCATAAACAAATCCACTTCTGC |
| SPE2 GE PCR R | *ADC2* | ATCTCACTTTACAGAAATTCTCGAA |
| SALK_073977 F | *ADC2* | ATAACCTTCTCACGTTGGTTC |
| SALK_073977 R | *ADC2* | GACAGGCCTCACATTCATC |
| LBb1.3 | *pROK2* (SALK) | ATTTTGCCGATTTCGGAAC |
| WiscDsLoxHs065_09H F | *ADC2* | TATCCGGTGAAATGTAACCA |
| WiscDsLoxHs065_09H R | *ADC2* | CACGATTCATCGTTTATTTATTTC |
| LB4 modified | *pDsLoxHs* (WiscDsLox) | GTAGATTTCCCGGACATGAAG |

**Supplemental Table S2. Plasmids used in transient assays and primers used for plasmid construction**

| Plasmid | Original plasmid | Construct | Primer (forward / reverse) |
|---|---|---|---|
| pKM5 | - | 35S::*Rluc* | – |
| pKN17 | pKM5 | 35S::s5′UTR:*Rluc* | RLUC 5UTR Fw / RLUC 5UTR Rv |
| pKM56 | - | 35S::*Eluc*-PEST | – |
| pYH21 | pKM56 | 35S::*ADC2*(WT):*Eluc*-PEST | SPE2 F / SPE2 R |
| pYH22 | pKM56 | 35S::*ADC2*(ΔCUG):*Eluc*-PEST | SPE2 d F / SPE2 d R |
| pYH40 | pKM56 | 35S::*ADC2*(ΔAUU):*Eluc*-PEST | SPE2 dAUU F / SPE2 dAUU R |
| pYH41 | pKM56 | 35S::*ADC2*(ΔAUU ΔCUG):*Eluc*-PEST | SPE2 dAUU F /SPE2 dAUU R |
| pYH65 | pKM56 | 35S::*ADC2*(ΔAUG):*Eluc*-PEST | SPE2 duORF F / SPE2 duORF R |
| pYH72 | pKM56 | 35S::*ADC2*(unpaired):*Eluc*-PEST | ADC-box unpaired F / ADC-box unpaired R |
| pYH73 | pKM56 | 35S::*ADC2*(paired):*Eluc*-PEST | ADC-box paired F / ADC-box paired R |

**Supplemental Table S3. Plasmids used for generation of transgenic or genome-edited lines and oligo DNA used for plasmid construction**

| Plasmid | Vector | Modification / Inserted component | Primer / oligo DNA |
|---|---|---|---|
| pYH74 | pKI1.1R | Deletion of Cas9/gRNA cassette and insertion of MCS | GG to CC IF for, GG to CC IF rev, CBSAE for, CBSAE for2 new, CBSAE rev |
| pYH75 | pYH74 | 35S::*ADC2*(WT):*Eluc*-PEST | - |
| pYH76 | pYH74 | 35S::*MASP1*(WT):*Eluc*-PEST | - |
| pYH70 | pKI1.1R | *SPE2* gR1 | SPE2 gR1 F, SPE2 gR1 R |
| pYH71 | pKI1.1R | *SPE2* gR2 | SPE2 gR2 F, SPE2 gR2 R |

**Supplemental Table S4. Primers used for plasmid construction**

| Name | Primer sequence (5′ to 3′) |
|---|---|
| RLUC 5UTR Fw | CTAGACTACAACTTCGCACCTTACCAAAAATGGCG |
| RLUC 5UTR Rv | TCGACGCCATTTTTGGTAAGGTGCGAAGTTGTAGT |
| SPE2 F | AGAACACGGGGGACTCTAGAACGCCTTCTCTTCTTCTTCTTC |
| SPE2 R | TCTTCTCTCTCTCCGTCGACGGCATCTTTATCTTCACCCTC |
| SPE2 d F | GATTCATCTTCTCTCTTCTCCTGA |
| SPE2 d R | TCAGGAGAAGAGAGAAGATGAATC |
| SPE2 dAUU F | GCGGGGATAGGTTCATCTTC |
| SPE2 dAUU R | GAAGATGAACCTATCCCCGC |
| SPE2 duORF F | TTCTGCAAAGTTGGGAGA |
| SPE2 duORF R | TCTCCCAACTTTGCAGAA |
| ADC-box unpaired F | CGGTTTTTAAAGGGGTACCTTTACAAATTCTGGGAA |
| ADC-box unpaired R | GGTACCCCTTTAAAAACCGGGCTCCGCGGAGCCTCGGCTACCC |
| ADC-box paired F | ACCCCGTAGGGCTCCGTCCGCGGAGCCCGGTTTTTAAAG |
| ADC-box unpaired R | GGTACCCCTTTAAAAACCGGGCTCCGCGGAGCCTCGGCTACCC |
| GG to CC IF for | ACTAGTGCGGCCGCCTGCACCTCGACCATATGGGAGA |
| GG to CC IF rev | AAACACTGATAGTTTAAACTGAAGGCGGGAAACGACA |
| CBSAE for | GATCGATAATTCGCGGTACATCGATCCTGCAGGGGATCCGAATTC |
| CBSAE for2 new | CTGCAGGGGATCCGAATTCCCGGGACCCAGCTTTCTTGTACAAAG |
| CBSAE rev | AGTTAGAATCTCAGGCCTTGTTGATATGAGGATTCGTG |

**Supplemental Table S5. Oligo DNA used for guide RNA cloning**

| Name | Sequence (5′ to 3′) |
|---|---|
| SPE2 gR1 F | ATTGGAGCCTCGGCTACCCCCTC |
| SPE2 gR1 R | AAACGAGGGGGTAGCCGAGGCTC |
| SPE2 gR2 F | ATTGTCTCCTGAGGGGGTAGCCG |
| SPE2 gR2 R | AAACCGGCTACCCCCTCAGGAGA |

**Supplemental Table S6. Primers used for RT-qPCR**

| Name | Target gene | Primer sequence (5′ to 3′) |
|---|---|---|
| ELUC qPCR for1 | *Eluc* | GAGAGCCTGCACAACTTCA |
| ELUC rev3 | *Eluc* | GTTCCTGTGGGTCTGCAT |
| SPE2 qPCR F | *ADC2* | CGAAGGTGAAAATGAAGAGGA |
| SPE2 qPCR R | *ADC2* | TCGCAGTAGTAAAACCCAAGG |
| EF1a qPCR for | *EF1α* | CCTTGGTGTCAAGCAGATGA |
| EF1a qPCR rev | *EF1α* | GAAGACACCTCCTTGATGATTT |
